# A genome-wide screen identifies genes in rhizosphere-associated *Pseudomonas* required to evade plant defenses

**DOI:** 10.1101/375568

**Authors:** Zhexian Liu, Polina Beskrovnaya, Ryan A. Melnyk, Sarzana S. Hossain, Sophie Khorasani, Lucy R. O’Sullivan, Christina L. Wiesmann, Jen Bush, Joël D. Richard, Cara H. Haney

## Abstract

*Pseudomonas fluorescens* and related plant root- (“rhizosphere”) associated species contribute to plant health by modulating defenses and facilitating nutrient uptake. To identify bacterial fitness determinants in the rhizosphere of the model plant *Arabidopsis thaliana*, we performed a Tn-Seq screen using the biocontrol and growth-promoting strain *Pseudomonas* sp. WCS365. The screen, which was performed in parallel on wild-type and an immunocompromised *Arabidopsis*, identified 231 genes that positively affect fitness in the rhizosphere of wild-type plants. A subset of these genes negatively affect fitness in the rhizosphere of immunocompromised plants. We postulated that these genes might be involved in avoiding plant defenses and verified 7 *Pseudomonas* sp. WCS365 candidate genes by generating clean deletions. We found that two of these deletion strains, Δ*morA* (encodes a putative diguanylate cyclase/phosphodiesterase) and Δ*spuC* (encodes a putrescine aminotransferase) formed enhanced biofilms and inhibited plant growth. Inhibition of plant growth by Δ*spuC* and Δ*morA* was the result of pattern triggered immunity (PTI) as measured by induction of an *Arabidopsis* PTI reporter and *FLS2/BAK1*-dependent inhibition of plant growth. We found that MorA acts as a phosphodiesterase to inhibit biofilm formation suggesting a possible role in biofilm dispersal. We found that both putrescine and its precursor arginine promote biofilm formation that is enhanced in the Δ*spuC* mutant, which cannot break down putrescine suggesting that putrescine might serve as a signaling molecule in the rhizosphere. Collectively, this work identified novel bacterial factors required to evade plant defenses in the rhizosphere.

**Importance:** While rhizosphere bacteria hold the potential to improve plant health and fitness, little is known about the bacterial genes required to evade host immunity. Using a model system consisting of *Arabidopsis* and a beneficial *Pseudomonas* sp. isolate, we identified bacterial genes required for both rhizosphere fitness and for evading host immune responses. This work advances our understanding of how evasion of host defenses contributes to survival in the rhizosphere.

## Introduction

Plant root-associated commensal microbes confer fitness advantages to plant hosts including growth promotion, nutrient uptake, and resistance to pathogens (1). In order to benefit its plant host, a root-associated microbe must survive in the rhizosphere, compete for plant nutrients, and avoid plant defenses. Despite the importance of rhizosphere competence for microbes to confer important benefits to plants, the mechanisms regulating rhizosphere fitness and evasion of host defenses are poorly understood.

Symbiotic bacteria must cope with a host immune system, which can robustly recognize microbe-associated molecular patterns (MAMPs) such as flagellin, lipopolysaccharide and chitin, and trigger a defense response. Model plant-associated *Pseudomonas* spp. can suppress some local plant defenses (2) and also trigger expression of MAMP-inducible genes (3). Using forward genetic approaches, genes required for rhizosphere competence have been identified in the model plant-associated bacterium, *Pseudomonas fluorescens* (4–7). Large-scale screens for *Pseudomonas* spp. fitness determinants in the plant rhizosphere have identified genes required for motility or nutrient uptake (4, 5). However, these factors have not been linked to evasion of plant immunity. How beneficial bacteria navigate the presence of a host surveillance system and manage to survive despite host defense responses remains elusive.

To identify fitness determinants in the presence of a plant immune system, we performed transposon mutagenesis coupled with high-throughput sequencing (Tn-Seq) using *Pseudomonas* sp. WCS365, on wild-type and immunocompromised *Arabidopsis* (see below). *Pseudomonas* sp. WCS365 is a growth promotion and biocontrol strain (6, 8, 9) and has been used for identification of *Pseudomonas* biofilm factors *in vitro* (10) and for genes important in rhizosphere colonization (6). Tn-seq is a high-throughput technique to rapidly assess fitness of each gene in a bacterial genome in a single experiment (11) and has been a particularly powerful method to identify determinants of bacterial fitness in association with animals (12–14) and plants (4). We reasoned that Tn-Seq might be an efficient method to rapidly identify bacterial genes required to avoid or suppress host immunity in the rhizosphere.

Here we report a Tn-Seq screen that identified 231 genes required for *Pseudomonas* sp. WCS365 fitness in the rhizosphere of wild-type *Arabidopsis* plants. We followed up on a subset of candidate genes that positively regulate fitness in the wild-type Col-0 rhizosphere but negatively affect fitness in the rhizosphere of a quadruple hormone mutant containing mutations in *DDE2, EIN2, PAD4*, and *SID2* [*deps*; (15)]. We found that two mutants, Δ*spuC* and Δ*morA*, induced pattern triggered immunity (PTI) in *Arabidopsis* via the flagellin receptor *FLS2*. We provide evidence that *Pseudomonas* sp. WCS365 *morA* and *spuC* temper biofilm formation, providing a possible link between bacterial physiology and induction of plant immunity.

## Results

### A Tn-seq screen identifies *Pseudomonas* sp. WCS365 fitness determinants in the *Arabidopsis* rhizosphere

To identify genes required for *Pseudomonas* sp. WCS365 fitness in the *Arabidopsis* rhizosphere, we performed a large-scale *mariner* transposon mutagenesis screen followed by next-generation sequencing (Tn-Seq) (see Methods for details). For this screen, germ-free *Arabidopsis* plants were grown in a sterile calcine clay and perlite mix with a plant nutrient solution (no carbon) to support plant growth. The screen was performed in parallel on wild-type *Arabidopsis* Col-0 and a mutant impaired in multiple hormone signaling pathways [*dde2-1/ein2-1/pad4-1/sid2-2; “deps”* mutant; (15)]. The *deps* mutant was chosen because it exhibited a 5- to 10-fold higher growth of *Pseudomonas* sp. WCS365 in the rhizosphere than wild-type plants (Fig. S1A). No-plant controls were supplemented with 20 mM succinate to support bacterial growth (Fig. S1B-C). We reasoned that insertions in WCS365 genes required for evasion of plant immunity would result in decreased fitness on wild-type plants but not on the *deps* mutant, allowing us to distinguish general colonization determinants from genes required to avoid or suppress plant immunity.

We sequenced the genome of *Pseudomonas* sp. WCS365 (Methods) to facilitate identification of transposon insertion sites in our Tn-Seq library (Genbank Accession PHHS01000000). To determine placement within the genus *Pseudomonas*, we generated a phylogenomic tree using 381 housekeeping genes identified by PhyloPhlAn [(16) Fig. S2]. We found that *Pseudomonas* sp. WCS365 falls within the *P. fluorescens* group of the fluorescent pseudomonads and is a close relative of *Pseudomonas* sp. NFM421within the *P. brassicacearum* subgroup (17).

To identify bacterial genes required for WCS365 fitness in the *Arabidopsis* rhizosphere, 3-week old plants were inoculated with a Tn-Seq library containing insertions in 66,894 TA dinucleotide sites distributed across the genome with approximately 9.8 insertions per 1000 bp (Fig. S3, Methods and Supplemental Data). Plants were inoculated with 10^4^ CFU per plant and plant roots or no plant controls were harvested one week later (Fig. 1A and Fig. S1B). We sequenced the transposon junctions in the rhizosphere and soil samples and compared the relative abundance of insertions in the rhizosphere of Col-0, the *deps* quadruple mutant, or the no plant control relative to the input. In our screen, we observed a significant bottleneck and ~35% of insertions were lost in any given treatment condition. Bottlenecks have previously been observed for other host-associated Tn-Seq screens (14). We adjusted our analysis to account for bottlenecks by first combining all reads per gene and then averaging the 3 replicates per gene (18).

**Figure 1.**
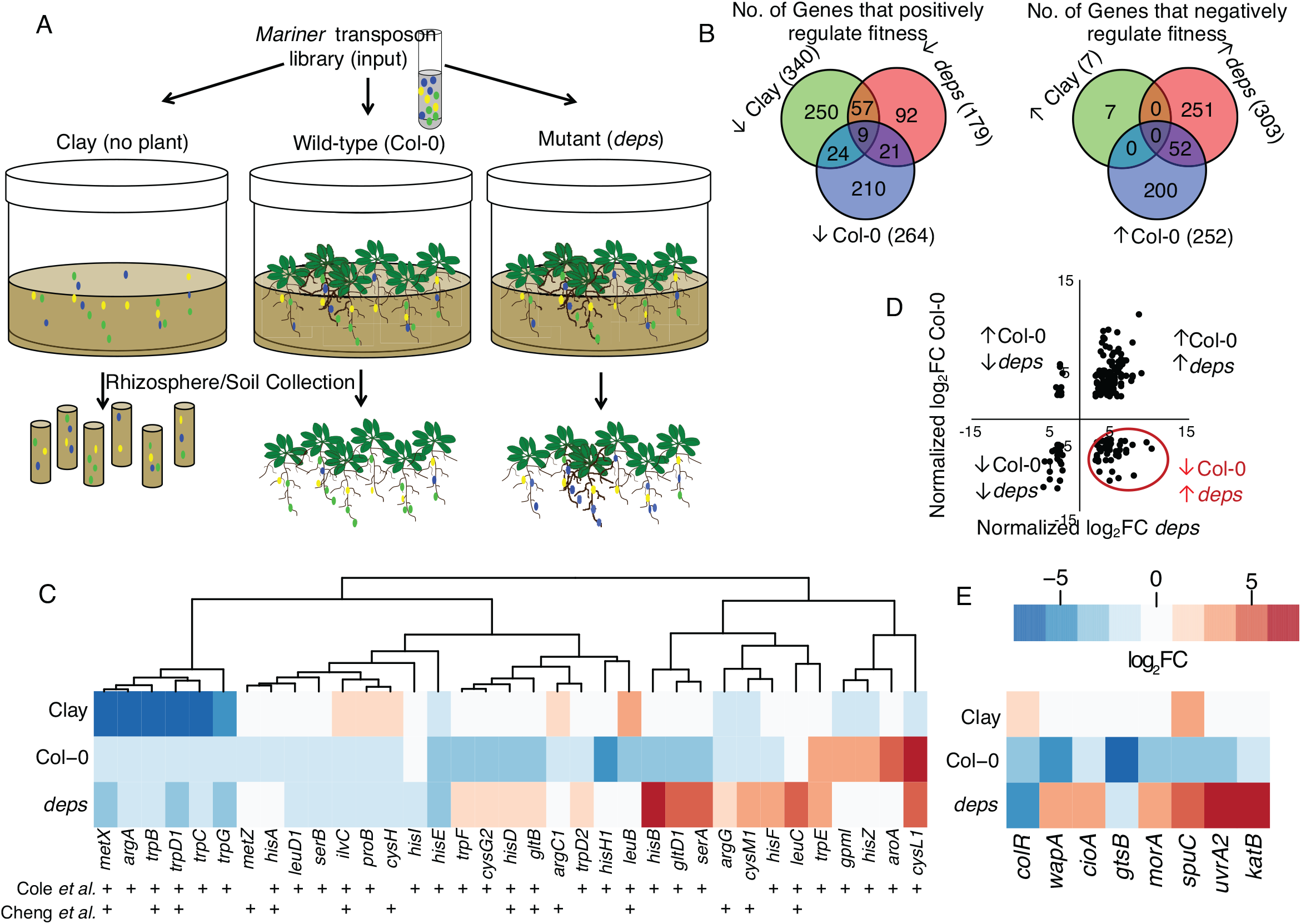
A Tn-Seq Screen identified *Pseudomonas* sp. WCS365 genes that affect fitness in the rhizospheres of wild-type and immuno-compromised *Arabidopsis*. (A) A transposon library was added to sterile clay with no plants, the roots of wild-type Col-0 plants, or a hormone mutant *deps*. (B) The Tn-Seq screen identified genes that positively or negatively affect fitness in the rhizosphere of wild-type Col-0 plants, an immuno-compromised hormone mutant (*deps*). (C) Amino acid biosynthetic genes previously shown to have positive effects on fitness [Cole et al., (4)] or negative effects on growth promotion [Cheng et al., (5)] in rhizosphere-associated *Pseudomonas* spp. (D) We focused on genes with a fitness cost in the *deps* rhizosphere but that provided a fitness advantage in the Col-0 rhizosphere. (E) Candidate genes chosen for follow-up showed significant differences with fitness in the Col-0 versus *deps* rhizospheres (*colR* was chosen as a control).

We identified 231 genes that positively affected fitness in the wild-type Col-0 rhizosphere (insertions in these genes caused a decrease in relative fitness; Fig. 1B and Supplemental Data). We found an additional 113 genes that positively affected fitness in the *deps* rhizosphere, but only 21 genes that positively affected fitness in the rhizosphere of both plants. We also found genes that negatively affected fitness in the rhizosphere of wild-type and the *deps* mutant (insertions in these genes enhanced relative fitness in the rhizosphere) including 52 genes that negatively affected fitness on both plant genotypes.

We compared the genes identified in our Tn-Seq screen to those identified in several recent screens for genes that affect the fitness or growth promotion ability of rhizosphere-associated *Pseudomonas* spp. (4, 5). Cole et al. (2017) found that insertions in amino acid biosynthesis genes resulted in a fitness advantage of *Pseudomonas* sp. WCS417 in the rhizosphere (4), while Cheng et al. (2017) found that insertions in orthologs of the WCS417 amino acid biosynthesis genes rendered *Pseudomonas* sp. SS101 unable to promote plant growth or protect from pathogens (5). We specifically looked at this same set of amino acid biosynthesis genes in our dataset and found that the majority of insertions in *Pseudomonas* sp. WCS365 amino acid biosynthesis genes reduced rhizosphere fitness in our study (Fig. 1C). Of note, a significant portion of insertions in these genes enhanced rhizosphere fitness in the *deps* mutant background (Fig. 1C). These results indicate that inability to synthesize certain amino acids results in a fitness defect in the wild-type Col-0 rhizosphere under the conditions in our study. These data also suggest that there may be altered amino acid profiles between the rhizosphere of wild-type plants and the *deps* mutant.

### *Pseudomonas* sp. WCS365 mutants have fitness defects in the rhizosphere

We hypothesized that bacterial genes that provide a fitness advantage in the presence of plant defenses might confer a fitness disadvantage in the absence of defenses responses. As a result, we considered genes that had a large negative log2fold-change ratio for fitness on Col-0 versus the *deps* mutant. We found that a significant portion of the genes that positively affected fitness in the Col-0 rhizosphere had negative effects on bacterial fitness in the *deps* rhizosphere (Fig. 1D-E and Supplemental Data). To determine if we successfully identified genes involved in survival of plant defenses, we retested 7 *Pseudomonas* sp. WCS365 candidates that were positive regulators of fitness in the Col-0 rhizosphere but negative regulators of fitness in the *deps* rhizosphere (Fig. 1E and Table 1). We generated clean deletions in: a catalase (*katB*), a diguanylate cyclase/phosphodiesterase (*morA*), a putrescine aminotransferase (*spuC*), an exinuclease (*uvrA*), a cytochrome oxidase subunit (*cioA*), an ABC transporter (*gtsB*), and a putative secreted protein (*wapA*) (Table 1; annotations were performed as described in the Methods). The previously identified colonization factor *colR* (7) was found to have impaired fitness in the rhizosphere of both wild-type plants and the *deps* mutant (Fig. 1E and Table 1), and was deleted as a control.

**Table 1.**
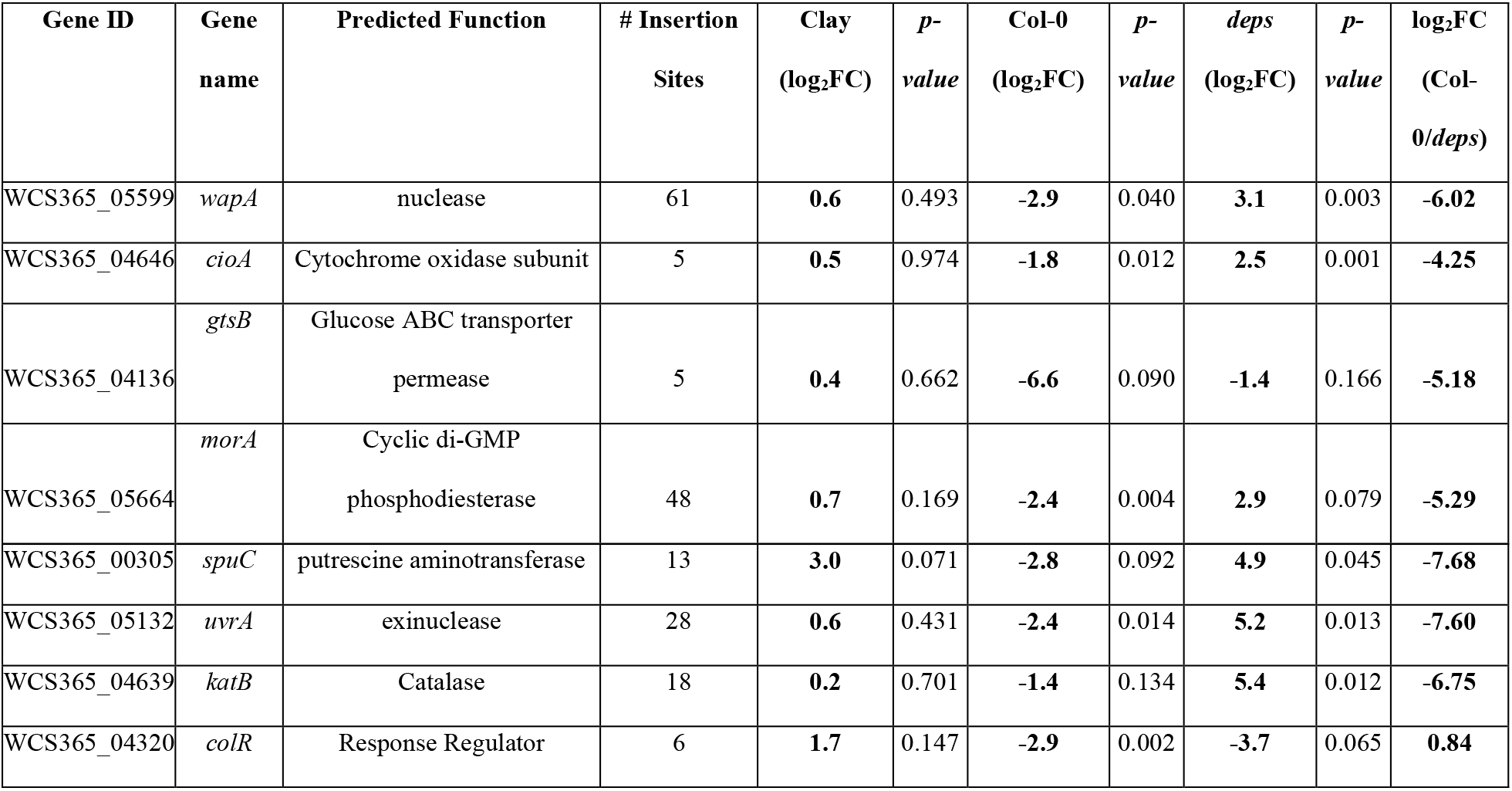
Genes selected for further characterization include those with a large different in fitness in the rhizospheres of WT (Col-0) and quadruple hormone mutant (*deps*) rhizospheres. Log_2_FC of Col-0/*deps* was calculated by first normalizing fitness data to the fitness in clay and then taking the ratio.

| Gene ID | Gene name | Predicted Function | # Insertion Sites | Clay (log<sub>2</sub>FC) | *p*-value | Col-0 (log<sub>2</sub>FC) | *p*-value | *deps* (log<sub>2</sub>FC) | *p*-value | log<sub>2</sub>FC (Col-0/*deps*) |
| --- | --- | --- | --- | --- | --- | --- | --- | --- | --- | --- |
| WCS365_05599 | *wapA* | nuclease | 61 | 0.6 | 0.493 | -2.9 | 0.040 | 3.1 | 0.003 | -6.02 |
| WCS365_04646 | *cioA* | Cytochrome oxidase subunit | 5 | 0.5 | 0.974 | -1.8 | 0.012 | 2.5 | 0.001 | -4.25 |
| WCS365_04136 | *gtsB* | Glucose ABC transporter permease | 5 | 0.4 | 0.662 | -6.6 | 0.090 | -1.4 | 0.166 | -5.18 |
| WCS365_05664 | *morA* | Cyclic di-GMP phosphodiesterase | 48 | 0.7 | 0.169 | -2.4 | 0.004 | 2.9 | 0.079 | -5.29 |
| WCS365_00305 | *spuC* | putrescine aminotransferase | 13 | **3.0** | 0.071 | **-2.8** | 0.092 | **4.9** | 0.045 | **-7.68** |
| WCS365_05132 | *uvrA* | exinuclease | 28 | **0.6** | 0.431 | **-2.4** | 0.014 | **5.2** | 0.013 | **-7.60** |
| WCS365_04639 | *katB* | Catalase | 18 | **0.2** | 0.701 | **-1.4** | 0.134 | **5.4** | 0.012 | **-6.75** |
| WCS365_04320 | *colR* | Response Regulator | 6 | **1.7** | 0.147 | **-2.9** | 0.002 | **-3.7** | 0.065 | **0.84** |

We retested our 7 *Pseudomonas* sp. WCS365 deletion mutants for growth and fitness in the *Arabidopsis* rhizosphere using a previously described hydroponic assay (9). We co-treated plants with wild-type *Pseudomonas* sp. WCS365 expressing mCherry from a plasmid and the *Pseudomonas* sp. WCS365 deletion strains expressing GFP from the same plasmid backbone.

We then quantified relative fluorescence as a measure of fitness. Under these conditions all 7 strains along with the Δ*colR* control had significant rhizosphere fitness defects (Fig. 2A). We tested the strains individually for colonization and found that a subset had significant growth defects in the rhizosphere (Fig. 2B). Collectively these results indicate that the Tn-Seq screen successfully identified novel *Pseudomonas* sp. WCS365 genes required for fitness in the *Arabidopsis* rhizosphere.

**Figure 2.**
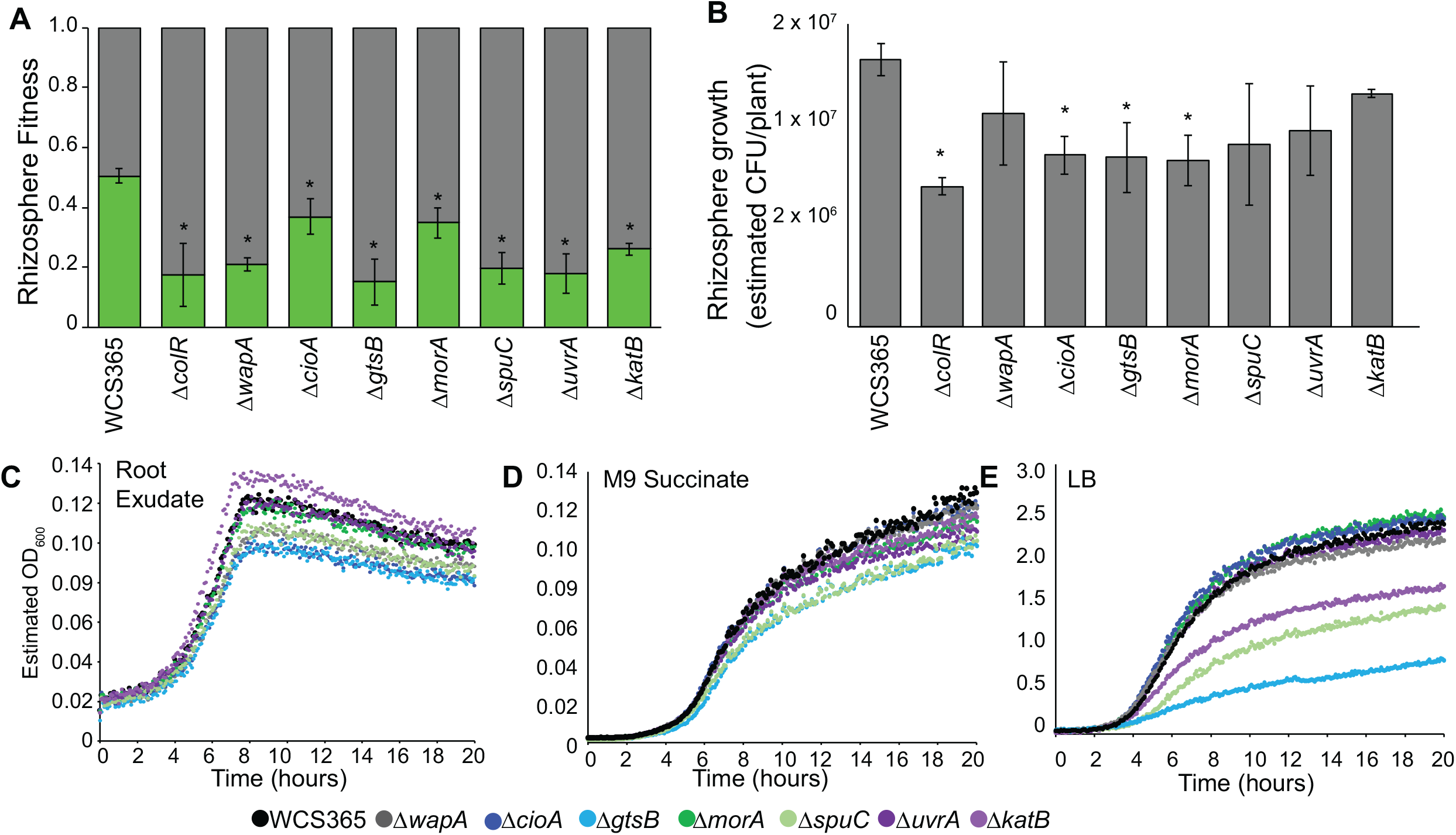
The Tn-seq screen identified *Pseudomonas* sp. WCS365 genes required for rhizosphere fitness. (A-B) Clean deletions were generated in *Pseudomonas* sp. WCS365 candidate genes and tested for fitness (A) or growth (B) in the *Arabidopsis* rhizosphere. Plants were grown in 48-well plates in hydroponic media. For fitness assays (A), plants were co-colonized with a wild-type strain expressing mCherry and a mutant or wild-type strain expressing GFP. The relative abundance was read after 3 days. For rhizosphere growth assays (B) plants were colonized with mutant or wild-type strains expressing GFP and fluorescence was measured as a proxy for growth. \**p*<0.01 by ANOVA and Tukey’s HSD relative to wild-type *Pseudomonas* sp. WCS365. (C-D) Growth of mutants in competition with wild-type cells in (C) LB (D) M9 + 30 mM succinate or (E) M9 + root exudate. Mutants expressing a GFP plasmid were growth in competition with wild-type expressing mCherry and growth of the GFP-expressing mutant was quantified with a plate reader. Quantification of growth rates and relative fitness are shown in Tables S1 and S2.

A previous screen for *Pseudomonas* fitness determinants in the rhizosphere identified mutants with poor or no growth in both minimal media and the rhizosphere (5). To determine if general growth defects underlie the observed rhizosphere fitness defects, we performed growth curves with root exudate as a sole carbon source and measured *in vitro* bacterial growth and fitness. We found that the majority of mutants showed normal growth rates (as measured by doubling time) when grown alone or in competition with wild-type (GFP mutant and mCherry wild-type) in root exudate (Fig. 2C and Fig. S4A). We found that a subset of the mutants had significant fitness defects in LB or minimal media as quantified by the fraction of the final culture that was composed of the mutant strain (Fig. 2C-E, Fig. S4A-C, and Tables S1 and S2). Because all strains could grow with root exudate a sole carbon source, these data suggest that fitness defects are specific to the presence of a live plant, or may be related to non-carbon related rhizosphere nutrients.

To gain insights into the requirements of these 7 mutants in the plant rhizosphere, we surveyed a publicly available fitness database where barcoded transposon libraries were assessed for fitness under *in vitro* conditions (19). By querying the database using the amino acid sequences from *Pseudomonas* sp. WCS365, we identified the loci with the highest similarity to the WCS365 predicted protein sequences the two most closely related strains, *Pseudomonas* sp. FW300-N2E2 and FW300-N2C3 (Supplementary Data). We found that insertions in N2E2 and N2C3 *morA, wapA*, and *katB* were fitness neutral under all conditions tested (Supplementary Data). Insertions in *gtsB* resulted in pleotropic fitness defects including during growth with glucose and galactose as sole carbon sources. Insertions in *spuC* resulted in growth defects with putrescine as a sole carbon or nitrogen source supporting a potential role in putrescine metabolism. Insertions in *uvrA* resulted in fitness defects in the presence of DNA-damaging agents supporting a role in DNA repair. Collectively, these data suggest that loss of these genes (with the exception of *gtsB*) do not have pleotropic growth defects, but rather defects that are specific to a limited number of conditions that may be relevant for rhizosphere growth.

While the majority of the 7 deletion mutants did not show growth or fitness defects *in vitro*, deletion of a predicted glucose transporter *gtsB* resulted in impaired growth rate and fitness of *Pseudomonas* sp. WCS365 in LB media (Fig. 2E, Tables S1 and S2). In the event the defect was due to a second site mutation, we reconstructed the Δ*gtsB* strain and independently confirmed the growth and fitness defect in LB. To test if *Pseudomonas* sp. WCS365 *gtsB* has a role in glucose transport, we tested the Δ*gtsB* mutant for growth in minimal media with succinate or dextrose as the sole carbon source. We found that the Δ*gtsB* mutant has a significant growth defect on dextrose but not succinate (Fig. S4D-E) consistent with its predicted role as a glucose transporter. Because glucose is not the dominant carbon source in LB media, it is unclear why this mutant would have a fitness defect in LB. As a result, it is unclear whether the Δ*gtsB* mutant fails in the rhizosphere due to an inability to transport glucose or due to pleotropic effects of deletion of this transporter component.

### *Pseudomonas* sp. WCS365 Δ*morA* and Δ*spuC* mutants induce pattern triggered immunity

When applied to wild-type *Arabidopsis thaliana* Col-0, *Pseudomonas* sp. WCS365 promotes plant growth as measured by increased plant weight and increased density of lateral roots (9). Microbe-associated molecular patterns (MAMPs) can be sensed by plants including *Arabidopsis* via interaction with pattern recognition receptors (PRRs) resulting in defense responses including callose deposition, inhibition of primary root growth, and induction of defense-related gene expression, collectively called pattern-triggered immunity (PTI) (2, 3). We therefore hypothesized that if any of the *Pseudomonas* sp. WCS365 genes identified in our screen are required to evade or suppress immunity, the deletion mutants might trigger PTI as measured by plant growth inhibition and induction of defense-related genes.

Under conditions where wild-type WCS365 promotes plant growth, we found that two of the seven mutants, Δ*morA* and Δ*spuC*, inhibited plant growth as measured by a reduction in plant lateral root density and primary root elongation (Fig. 3A-B; Fig. S5). The remaining mutants including the Δ*gtsB* mutant, which had the most severe rhizosphere growth and fitness defect (Fig. 2A-B), still triggered an increase in *Arabidopsis* lateral root density. We tested an *Arabidopsis* reporter line consisting of the promoter of MAMP-inducible gene *MYB51* fused to the ß-glucuronidase (*MYB51pro::GUS*) reporter gene, which provides a qualitative readout of PTI (2). We found slight induction of *MYB51* by wild-type *Pseudomonas* sp. WCS365 and enhanced *MYB51* expression in seedlings exposed to Δ*morA*, Δ*spuC*, or flg22 (Fig. 3C). Collectively, these data suggest that *morA* and *spuC* are required to avoid triggering PTI in *Arabidopsis*.

**Figure 3.**
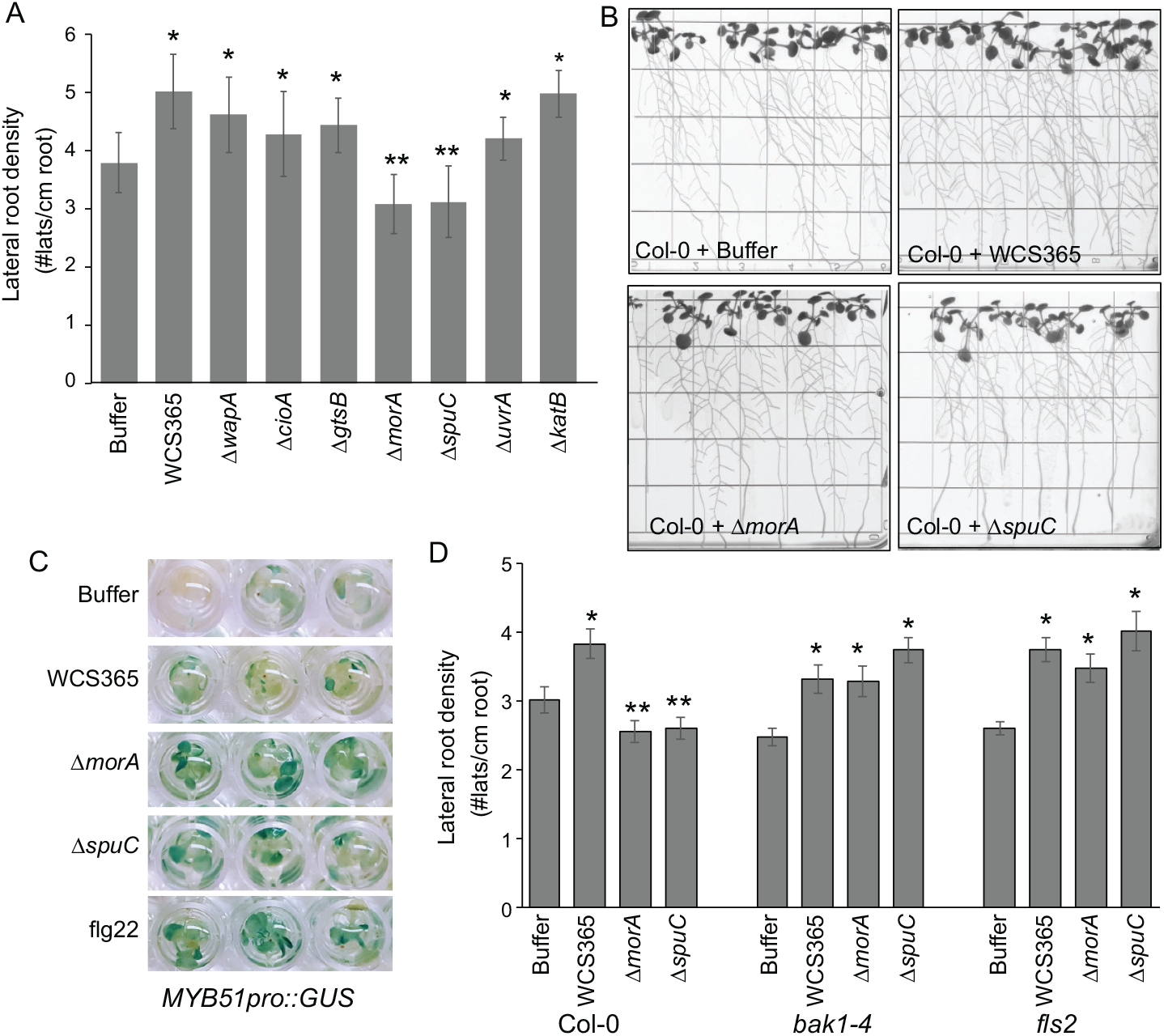
*Pseudomonas* sp. WCS365 Δ*morA* and Δ*spuC* mutants induce Pattern Triggered Immunity. (A) The growth promotion ability of *Pseudomonas* sp. WCS365 mutants was tested on wild-type *Arabidopsis thaliana* ecotype Col-0. Lateral root density (lateral roots per cm of primary root) is shown. (B) Images of growth promotion assays showing the Δ*morA* and Δ*spuC* mutants. PGP assays with the remainder of *Pseudomonas* sp. WCS365 mutants are shown in Fig. S5. (C) Using an *Arabidopsis* MAMP inducible transgenic reporter line *(MYB51pro::GUS)*, we found that *Pseudomonas* sp. WCS365 Δ*morA* and Δ*spuC* mutants induce MAMP-dependent gene expression. (D) *Arabidopsis* growth inhibition by the Δ*morA* and Δ*spuC* mutants is dependent on MAMP perception via *BAK1* and *FLS2*. \**p*<0.05 increase in lateral root density; \*\**p*<0.05 decrease in lateral root density relative to Buffer-treated controls by ANOVA and Tukey HSD.

Plant perception of the majority of MAMPs in *Arabidopsis* is dependent on the co-receptor BAK1 (20). We therefore tested if growth promotion by Δ*morA* and Δ*spuC* is restored in a *bak1-4* mutant. We observed significant growth promotion of an *Arabidopsis bak1-4* mutant by *Pseudomonas* sp. WCS365 Δ*morA* and Δ*spuC* (Fig. 3D). These data indicate that the Δ*morA* and Δ*spuC* inhibit plant growth due to induction of PTI via *BAK1*.

Motility is necessary for rhizosphere colonization by a number of microbes (21); however, failure to downregulate motility might cause induction of PTI as *Arabidopsis* can sense flagellin produced by *Pseudomonas* spp. We tested if growth inhibition was dependent on the plant flagellin perception by testing an Arabidopsis *FLS2* mutant that cannot sense bacterial flagellin (22). We observed significant growth promotion of an *Arabidopsis fls2* mutant by *Pseudomonas* sp. WCS365 Δ*morA* and Δ*spuC* (Fig. 3D). These data indicate that *Arabidopsis* flagellin perception underlies the defense response triggered by the Δ*morA* and Δ*spuC* mutants.

### *Pseudomonas* sp. WCS365 Δ*morA* and Δ*spuC* mutants form enhanced biofilms without defects in motility

Diguanylate cyclases, including *P. aeruginosa* MorA, are positive regulators of biofilm formation and negative regulators of swimming motility (23, 24). As a result, we predicted that the Δ*morA* mutant would have increased motility and decreased biofilm formation and that this might explain its rhizosphere fitness defect and *FLS2*-dependent inhibition of growth. Unexpectedly, we found that neither the Δ*morA* mutant nor the majority of other *Pseudomonas* sp. WCS365 deletion mutants had increased swimming motility (Fig. 4A and S6A). We found several mutants had subtle decreases in surfing motility [(25); Fig. S6B]. The deletion of the ABC transporter *gtsB* resulted in consistently impaired surfing and swimming motility (Fig. S6A-B). Collectively these data indicate that increased bacterial motility does not underlie the rhizosphere fitness defect or induction of defenses in these mutants.

**Figure 4.**
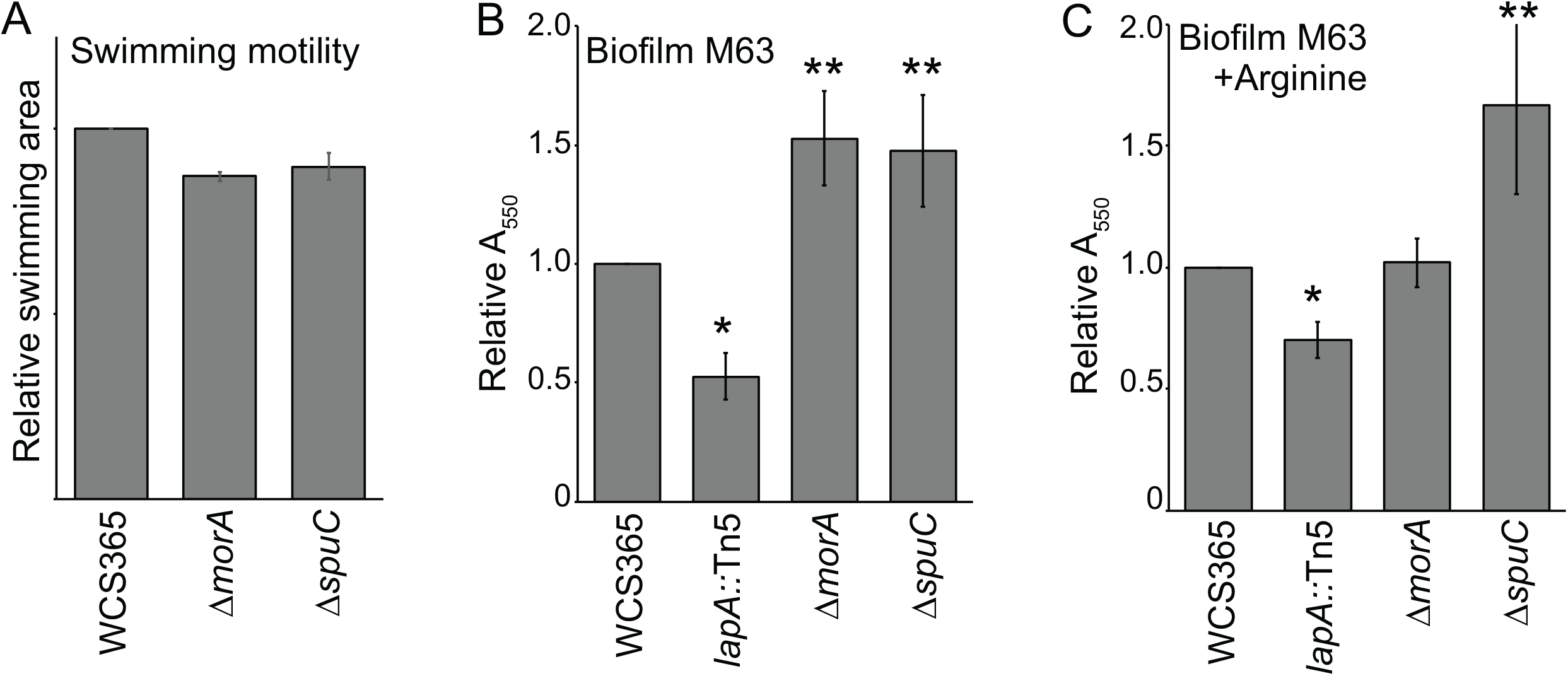
*Pseudomonas* sp. WCS365 Δ*spuC* and Δ*morA* mutants do not have motility defects and form enhanced biofilms. (A) Swimming motility and (B-C) crystal violate biofilm assays with *Pseudomonas* sp. WCS365 Δ*morA* and Δ*spuC* in (B) M63 media or (C) M63 media supplemented with arginine. The *lapA*::Tn5 mutant was used as a positive control for a biofilm-impaired mutant. Data are the average of 4-5 biological replicates with 3-4 technical replicates per experiment. \**p*<0.05 decrease in biofilm formation; \*\**p*<0.05 increase in biofilm relative to wild-type WCS365 by ANOVA and Tukey’s HSD.

MorA has both predicted diguanylate cyclase (DGC) and phosphodiesterase (PDE) domains. DGC domains are negative regulators of biofilm formation by promoting c-di-GMP accumulation, while PDEs decrease biofilm formation and promote dispersal by lowering c-di-GMP levels (26). As a result, we tested whether the Δ*morA* and the remaining WCS365 mutants had alterations in biofilm formation in a standard *in vitro* crystal violet assay in M63 minimal media or M63 salts supplemented with arginine, which has previously been shown to enhance biofilm formation in *Pseudomonas* spp. (27, 28). We found that both the Δ*morA* and Δ*spuC* mutants formed strongly enhanced biofilms in M63 media and Δ*uvrA* and Δ*wapA* formed weakly enhanced biofilm (Fig. 4B and S6C). Additionally, we found that only the Δ*spuC* mutant formed enhanced biofilms in M63 supplemented with arginine (Fig. 4C and S6D). SAD-51, a known surface attachment deficiency mutant of *Pseudomonas* sp. WCS365 with a transposon insertion in *lapA* (10), was used as a control. Enhanced biofilm formation by *Pseudomonas* sp. WCS365 Δ*morA* and Δ*spuC* mutants *in vitro* suggests that hyperbiofilm formation or inability to disperse may underlie their fitness defects in the rhizosphere.

### The phosphodiesterase activity of *Pseudomonas* sp. WCS365 MorA inhibits biofilm formation and is required for rhizosphere fitness

*Pseudomonas* sp. WCS365 *morA* encodes a putative diguanylate cyclase/phosphodiesterase A (DGC/PDEA) gene homologous to *P. aeruginosa* PAO1 *morA*, a known regulator of biofilms and virulence (23, 24). To determine if the phosphodiesterase activity or diguanylate cyclase activity are necessary for rhizosphere fitness, we generated point mutations in the GGDEF and EAL domains to inactivate the diguanylate cyclase (*morA^GGAAF^*) and phosphodiesterase domains (*morA^AAL^*) (Fig. 5A).

**Figure 5.**
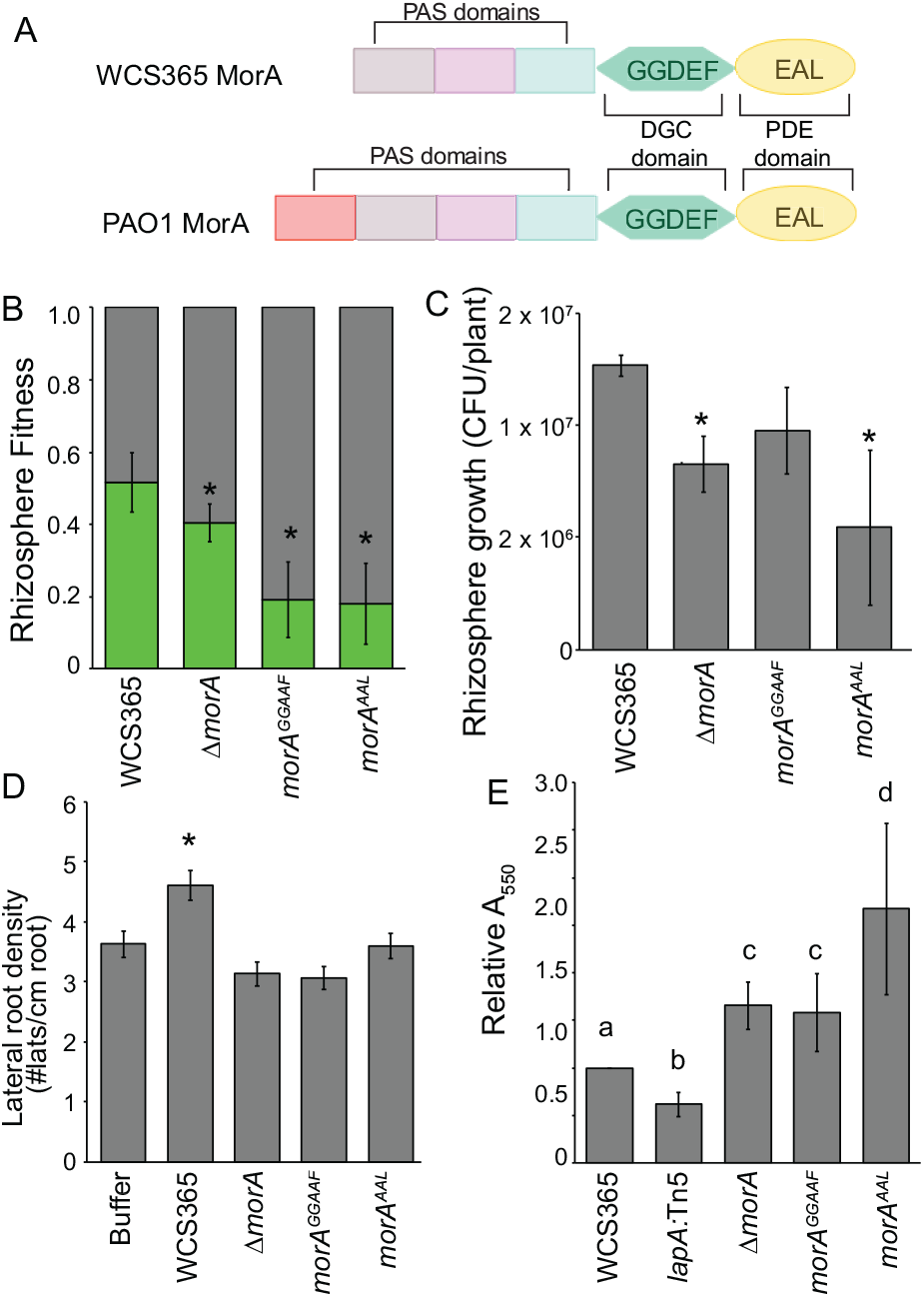
*Pseudomonas* sp. WCS365 *morA* acts as a phosphodiesterase to enhance rhizosphere fitness and negatively regulate biofilm formation. (A) Illustration of functional domains of MorA homolog in *Pseudomonas* sp. WCS365 and *P. aeruginosa* PAO1. *P. fluorescens* WCS365 MorA and *P. aeruginosa* PAO1 MorA share a similar predicted domain organization. *P. aeruginosa* PAO1 MorA contains an additional PAS-PAC sensor domain that contains a putative heme pocket. Point mutations in the predicted catalytic sites of the MorA diguanylate cyclase (*morA^GGAAF^*) and phosphodiesterase (*morA^AAL^*) domains were tested for (B) rhizosphere fitness, (C) rhizosphere growth, (D) growth promotion and (E) biofilm formation. (B-D) \**p*<0.01 by Student’s t-test; data are the average of 3 biological replicates with at least 8 plants per replicate. (E) letters designate levels of significance (*p*< 0.05) by ANOVA and Tukey’s HSD; data are the average of 4 biological replicates with 8 technical replicates per experiment.

We tested the *morA^GGAAF^* and *mor^AAL^* mutants for rhizosphere fitness, rhizosphere growth, and biofilm formation. Surprisingly we found both the *morA^GGAAF^* and *mor^AAL^* mutants retained defects in rhizosphere growth and fitness as well as plant growth promotion (Fig. 5B-D). We found the *morA^AAL^* mutant had even greater biofilm formation in a crystal violet assay and the *morA^GGAAF^* mutant retained the enhanced biofilm formation of the Δ*morA* mutant (Fig. 5E). That the *morA^GGAAF^* mutant retains the Δ*morA* phenotype suggests that the conserved GGDEF motif does not contribute to diguanylate cyclase activity. Collectively, these data suggest that *Pseudomonas* sp. WCS365 MorA acts as a phosphodiesterase to temper biofilm formation or promote dispersal in the rhizosphere.

### Putrescine acts as a signaling molecule to promote *Pseudomonas* sp. WCS365 biofilm formation

Putrescine is present in the rhizosphere of tomato (29) and so we wondered if putrescine could serve as a signaling molecule to promote bacterial biofilm formation in the rhizosphere. We found that the Δ*spuC* mutant formed significantly enhanced biofilms in the presence of arginine (Fig. 4C). Arginine can be converted to putrescine in *P. aeruginosa* PAO1 (30). In *P. aeruginosa*, SpuC breaks down putrescine into 4-aminobutyraldehyde, which can be further broken down into succinate and used as a carbon source (30) and so in the presence of arginine, an *spuC* mutant should over-accumulate putrescine. We tested whether the *Pseudomonas* sp. WCS365 Δ*spuC* was impaired in the catabolism of putrescine by testing whether the Δ*spuC* mutant could use putrescine as a sole carbon source. While wild-type *Pseudomonas* sp. WCS365 could grow in minimal media with 25 mM putrescine as the sole carbon source, the Δ*spuC* mutant was severely impaired (Fig. S7), indicating that this mutant does indeed fail to metabolize putrescine.

If putrescine is serving as a signaling molecule in the rhizosphere, we reasoned that other genes involved in putrescine synthesis or metabolism should also have fitness defects in our experiment. We reconstructed the putrescine uptake, synthesis and utilization pathways based on what is known in WCS365 and other organisms (30–32) (Fig. 6A). We identified putrescine uptake system operon *potFGHI* (WCS_00300-WCS_00304), a gene involved in conversion of arginine to putrescine (*speA* WCS365_02314; *aguA* WCS365_01963; *aphA* WCS365_00490). Notably, we were unable to identify a homologue of *P. aeruginosa aguB* [acts with *aphA* to catalyzes the conversion of N-carbamoylputrescine to putrescine (33)] in the WCS365 genome or in the genome of the close relative *Pseudomonas* sp. NFM421. This indicates that either *Pseudomonas* sp. WCS365 cannot convert arginine to putrescine, that *aphA* alone catalyzes this reaction, or that a different enzyme substitutes for *aguB*. For putrescine conversion to succinate, we identified *pauC, gabT* and *gabT* homologues in WCS365 (WCS365_03989, WCS365_05732, and WCS365_05733).

**Figure 6.**
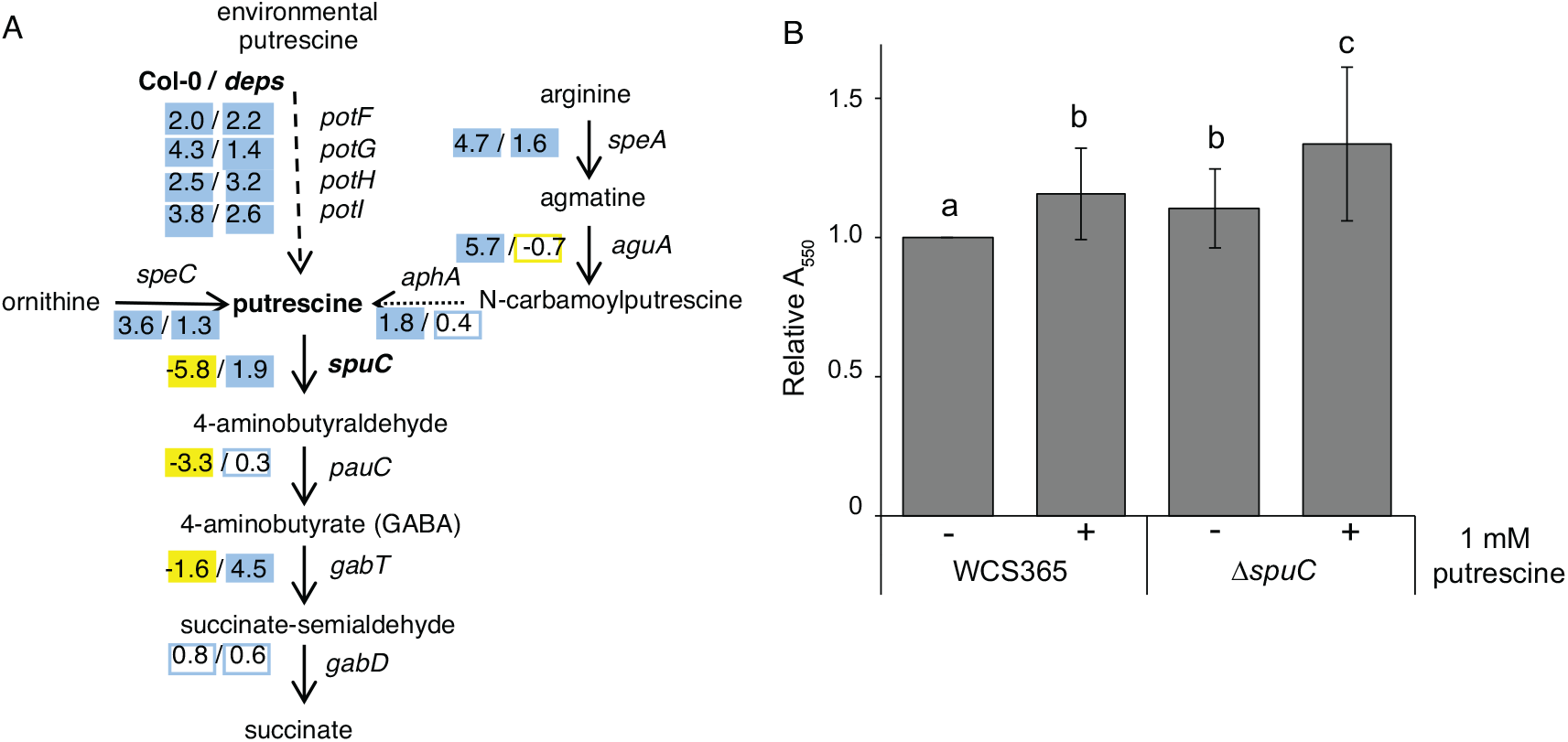
Putrescine promotes biofilm formation in *Pseudomonas* sp. WCS365. (A) Putrescine uptake, synthesis and metabolism pathway in *Pseudomonas* sp. WCS365 with log_2_FC fitness data from the Tn-Seq experiment shown. (B) Crystal violet assays were performed in M63 media or M63 media with 1 mM putrescine. Data shown are the average of 6 biological replicates with 8 technical replicates per experiment. Letters designate *p*<0.05 by ANOVA and t tests.

We overlaid our Tn-Seq fitness data onto the putrescine uptake, synthesis and utilization pathway and found that insertions in *spuC, pauC* and *gabT*, which are involved in the conversion of putrescine to succinate, all positively regulate fitness in the rhizosphere of wild-type but not immunocompromised plants (Fig. 6A). Interestingly, we found that all genes potentially involved in putrescine uptake or synthesis had increased fitness scores in our Tn-Seq experiment indicating that they are negative regulators of fitness (Fig. 6A). These data are inconsistent with putrescine being a significant carbon source in the rhizosphere; if it were, we would predict that a loss of uptake or synthesis would impair fitness in the rhizosphere. Rather they support a role for putrescine as a signaling molecule that promotes biofilm formation in the rhizosphere.

We tested whether putrescine could directly promote biofilm formation in wild-type *Pseudomonas* sp. WCS365 and the Δ*spuC* mutant and found that putrescine is sufficient to promote biofilm formation in wild-type bacteria (Fig. 6B). Furthermore, the putrescine-mediated biofilm enhancement is exacerbated in the Δ*spuC* mutant (Fig. 6B). These data support the hypothesis that putrescine may serve as a signaling molecule to trigger a *Pseudomonas* sp. WCS365 lifestyle change in the rhizosphere.

## Discussion

Here we report a screen that identified a *Pseudomonas fluorescens* WCS365 putrescine aminotransferase (SpuC) and a phosphodiesterase (MorA) that are required to evade plant defenses. Deletion of either gene results in induction of pattern triggered immunity (PTI) in *Arabidopsis* as measured by *FLS2/BAK1*-dependent inhibition of plant growth, and increased induction of the MAMP-inducible *MYB51* gene (Fig. 3). Previous studies have found that *Pseudomonas* spp. induce a subset of plant PTI responses while suppressing others (2, 3). Collectively these data reveal novel mechanisms used by *Pseudomonas* sp. to avoid detection by a plant host.

A previous study found that an insertion between *spuC* and the *potFGHI* uptake system in WCS365 increased putrescine uptake and decreased rhizosphere fitness (34). In *P. aeruginosa*, SpuC is involved in the catabolism of putrescine into 4-aminobutyraldehyde (30); as the *Pseudomonas* sp. WCS365 Δ*spuC* mutant cannot utilize putrescine (Fig. S7) it may accumulate putrescine or related compounds. We found that arginine and putrescine promote biofilm formation in wild-type *Pseudomonas* sp. WCS365, and that loss of the *spuC* gene further enhances the biofilm-promoting effects of arginine and putrescine (Fig. 4C and Fig. 6). Putrescine is a positive regulator of biofilm formation in *Yersinia pestis* (35); while it is possible that the fitness defect in the Δ*spuC* mutant is due to an inability to metabolize putrescine as a sole carbon source, we propose instead that putrescine serves as a signal that informs *Pseudomonas* sp. of the presence of a eukaryotic host such as a plant. This in turn triggers a bacterial lifestyle switch to promote attachment and biofilm formation. Loss of the *spuC* gene results in hypersensitivity to exogenous putrescine resulting in changes in bacterial physiology that ultimately triggers plant defenses. Collectively, these data suggest that either arginine or putrescine in the plant rhizosphere might act as a signaling molecule to trigger a lifestyle change and evade plant defenses.

By performing an in-depth characterization of the role of a *Pseudomonas* sp. WCS365 diguanylate cyclase/phosphodiesterase A (DGC/PDEA) gene, *morA*, in rhizosphere competence and biofilm formation, we determined that MorA primarily acts as a phosphodiesterase to temper biofilm formation in the rhizosphere (Fig. 5). Bacterial DGC/PDEA activities regulate the intracellular levels of the bacterial second messenger cyclic diguanylate (c-di-GMP). In *P. aeruginosa*, DGCs promote c-di-GMP synthesis and positively regulate biofilm formation while subsequent lowering of c-di-GMP by PDEAs downregulates biofilm production and promotes dispersal (26). Because *morA* acts as a PDEA, this suggests its role may be to temper biofilm formation and/or promote dispersal in the rhizosphere. Upregulation of flagellin biosynthesis also accompanies dispersal; however, the WCS365 *morA* mutant has no defect in swimming motility (Fig. 5A) indicating it can still produce flagellin. These data suggest that ability to disperse or downregulate biofilm production may be required to evade induction of host defenses.

The roles of the other 5 genes that we confirmed rhizosphere fitness defects for in our study (Fig. 2A; *katB, uvrA, wapA, cioA*, and *gtsB*) can be predicted from known functions in other organisms or data presented in our paper. The catalase KatB is necessary for detoxification of hydrogen peroxide in *P. aeruginosa* (36), which suggests that *Pseudomonas* sp. WCS365 must detoxify reactive oxygen to compete in the rhizosphere. *uvrA* in *Escherichia coli* encodes a subunit of the ABC excinuclease responsible for the repair of UV-induced thymine dimer and other DNA damage (37, 38); a requirement for *uvrA* in the rhizosphere suggests that *Pseudomonas* sp. might need to contend with DNA damage. The *cioA* gene is highly similar to the previously characterized *P. aeruginosa* PAO1 *cioA* (79% identity, 98% coverage), which, together with the gene product of *cioB*, forms CIO, a cytochrome with low affinity to oxygen (39) and might indicate a necessity to adapt to oxidative stress. A requirement for *gtsB* (PFLU4845), which has been shown to encode a glucose permease subunit of a ATP-binding cassette transporter (40), suggests that glucose may be a significant carbon source in the *Arabidopsis* rhizosphere. Interestingly, WapA toxins are associated with a cognate immunity protein WapI in *B. subtilis* 168 and are involved in cell-cell killing (41); however, no putative *wapI* gene was identified in the vicinity of *wapA* in *Pseudomonas* sp. WCS365 contigs, possibly due to the highly polymorphic sequences of WapA proteins. How loss of *wapA* leads to rhizosphere fitness defect remains to be elucidated. Collectively, the requirement of these genes in rhizosphere fitness implicates diverse bacterial pathways and processes in survival in the *Arabidopsis* rhizosphere.

In mixed inoculations, we observed that plant defenses were targeted only towards specific mutant bacteria and not the community as a whole. We used two mixed inoculation strategies: a Tn-Seq community and 1:1 inoculations of wild-type and mutant bacteria. With the Tn-Seq library, we observed similar levels of colonization by the library and wild-type bacteria, this indicates that any defense response that was effective at killing bacteria was highly localized. The second mixed inoculation strategy was using a 1:1 ratio of wild-type or mutant bacteria. Under these conditions, we did observe a slight but not statistically significant decrease in the total number of bacteria. This again indicates that any defense responses must be highly localized at the level of one or a group of cells rather than the entire plant. Collectively these data suggest a model where a highly localized defense response may help target a pathogen invader without disrupting the complex and diverse rhizosphere microbial community. As a result, we hypothesize that Δ*spuC* and Δ*morA* mutants are poor rhizosphere colonizers due to an inability to disperse from the initial site of colonization after triggering plant immune response.

In summary, evasion or suppression of the plant immune system is essential for pathogens to successfully infect their plant hosts. Our results support a growing body of evidence that avoiding plant defenses is also critical for survival of commensals in association with a host (42). Many studies point to attachment being critical for virulence of bacterial pathogens (43) and colonization of commensals (44). However, our work shows a positive correlation between hyperformation of biofilm and induction of plant defenses. This work indicates that changes in bacterial physiology may be necessary for evasion of plant defenses and survival in association with a eukaryotic host.

## Materials and methods

### Plant growth

For the Tn-Seq screen, seeds were sterilized using chlorine gas which successfully eliminates detectible endophytes by 16S rRNA sequencing (9). For growth promotion and root colonization experiments, seeds were sterilized using either chlorine gas (100 mL bleach + 3 mL concentrated hydrochloric acid in an air-tight container for 4 hours) bleach sterilization (70% ethanol for 2 minutes followed by 10% bleach for 5 minutes followed by three washes in sterile distilled water). Plants were grown under 16h/8h day/night 22°C at 100 μE light.

*Arabidopsis* genotypes were in the Col-0 background and included *dde2-1/ein2-1/pad4-1/sid2-2; “deps”* mutant (Tsuda et al. 2009), *bak1-4* (45), *fls2* (22), *MYB51pro-GUS* (2), *WRKY11pro-GUS* (2).

### Strains, media, and culture conditions

For routine culturing, *Pseudomonas* sp. WCS365 was grown on Luria-Bertani (LB) agar or in LB medium at 28°C; *Escherichia coli* strains were grown on LB agar or in LB medium at 37°C. Antibiotics were used at the following concentrations when appropriate: gentamicin 5 μg/mL (*E. coli*) or 10 μg/mL (*Pseudomonas*) or nalidixic acid 15 μg/mL.

### Genome sequencing of *Pseudomonas* sp. WCS365 and phylogenomic analysis

Bacteria DNA was isolated using Qiagen Purgene Kit A and sonicated into ~500 bp fragments. Library construction and genome assembly was performed as described (46, 47). The draft genome of *Pseudomonas* sp. was assembled into 60 contigs containing 6.56 Mb and a predicted 5,864 coding sequences. The WCS365 Whole Genome Shotgun project has been deposited at DDBJ/ENA/GenBank under the accession PHHS00000000. The version described in this paper is version PHHS01000000.

### Tn-Seq library preparation

A mariner transposon [pSAM_DGm; (48)] was introduced in *Pseudomonas* sp. WCS365 via conjugation with SM10λpir *E. coli*. We found the transposon integrated into the *E. coli* genome with detectible frequency and so we minimized culture growth time to ~6 hours. The conjugation was left at 28°C for 2 hours to minimize replication and the occurrence of sibling mutants. The conjugation was then scraped off of plates and frozen. The mating mix was plated on LB in 100 mM petri dishes with gentamicin (10 μg/ml) and nalidixic acid (15 μg/ml) at a density of 1500-3000 colonies per plate. After approximating the number of colonies, they were scraped off the plates and pooled to ~10,000 colonies (4-6 plates) per mix. This was repeated 10 independent times from 10 mating mixes for a total of an estimated 100,000 colonies. The OD_600_ of each independent pool was measured and to make the final pool, the density was normalized so the approximate same number of cells from each colony was added. The library was diluted to an OD_600_ = 0.2 and allowed to recover for 1 hr in LB prior to aliquotting and freezing. An aliquot of the library was plated and CFUs were counted before the final plant inoculation step.

We sequenced the region flanking the transposon insertions in our library (Fig. S3 and Methods). We mapped the insertion sites to the 6.56 Mb draft *Pseudomonas* sp. WCS365 genome and found that the library contained insertions in 66,894 TA dinucleotide sites distributed across the genome with approximately 9.8 insertions per 1000 bp. Of the 5,864 annotated genes in WCS365, we identified insertions in 5,045 genes. Distribution of insertions by gene is shown in Fig. S3B and the gene list in the Supplemental Data; we found a mean of 10.3 and median of 8 insertions per gene.

### Tn-Seq Experimental setup and sequencing methods

Sterile plant growth substrate was prepared by mixing 2 parts Turface Prochoice calcine clay, 2 parts Turface quickdry and 1 part perlite. The mixture was washed 10 times with distilled water to remove soluble nutrients. 12 cm diameter plant tissue culture vessels (C1775; Phytotechnology Laboratories) were filled 3 cm deep with the mixture ensuring the mixture was saturated with water but water did not pool in the box. 10 mL of Hoagland’s Solution was added to each box (Fig. S1). The end result was a porous growth substrate. Boxes were capped and autoclaved.

To facilitate *Arabidopsis* germination, twenty plugs of MS agar (1× with 2% sucrose) 3 mm^2^ in diameter were evenly distributed on the growth medium surface and seeds were sowed directly on the agar plugs (Fig. S1). 10 boxes per treatment were used with 20 plants per box (n = 200 plants) and three biological replicates were used per treatment. Treatments included Col-0, the *deps* mutant and no plant control (20 mM succinate was added on top of the plugs in lieu of plants as a bacterial nutrient source). Plants were grown for 2 weeks prior to inoculation. The library was diluted to 5 × 10^4^ CFU/mL based on plate counts and 200 μl was added to each plant or each control agar plug for a total of 10^4^ bacteria per plant. 20 plants or no-plant equivalents were inoculated per box for a total of 2 × 10^5^ bacteria per box and 2 × 10^6^ bacteria sampled for the entire experiment; each insertion was represented about 20 times in the original inoculum. We found that each plant could support a total of 5 × 10^7^ CFU/gram (Col-0) and that each plant weighed about 50 mg at the end of the experiment meaning our pool grew out 250 fold over the course of the experiment. Bacteria were allowed to grow for 1 week before harvesting.

To harvest bacteria, plants were removed from the growth substrate and loose soil was removed (Fig. S1). DNA was isolated using MoBio Power Soil DNA isolation kit (columns for up to 10 grams of material); all material from a single treatment and replicate was processed together. Yields were on the order of 3 to 12 μg of DNA from the plant and clay samples and 3 μg of DNA was used as an input for library construction.

Sequencing libraries were prepared as described using cleavage with the MmeI enzyme with modifications (49). Adapter and primer sequences along with a schematic of library construction can be found in Fig. S3. Sequencing libraries include 3 reps of 1) the input library, 2) Col-0 rhizosphere, 3) the hormone mutant (*dde2-2, ein2-1, pad4-1, sid2-2*) and 4) no plant treatments. Each replicate (12 samples total) was indexed separately. Sequencing libraries were prepared by digesting input DNA with the MmeI enzyme, end repairing, and ligating a double stranded blunt-ended adapter molecule. Transposon and adapter-specific primers were used to amplify the region flanking the transposon (the transposon is palindromic and so both directions should amplify with similar frequency). The presence of a predicted 169 bp product was confirmed with an Agilent Bioanalyzer. All twelve samples were pooled and run in the same Illumina HiSeq lane using single end 50 bp reads.

### Tn-Seq data analysis

Data analysis was performed using Galaxy and a modified version of the MaGenTA pipeline described in (18). Our custom adapters and barcodes are shown in Supplementary Data and a schematic of library construction is shown if Fig. S3. The adapter was trimmed using the custom sequence 5’ ACAGGTTGGATGATAAGTCCCCGGTCT 3’. Sequencing reads were trimmed to remove the transposon sequencing so 21-22 bp that represented the flanking region post MmeI cleavage remained. After barcode splitting and trimming, between 458,679 and 1,298,597 reads were assigned to each individual treatment. Sequences were mapped back to the *Pseudomonas* sp. WCS365 draft genome using Map_with_Bowtie_for_Illumina using the following custom settings: −n = 1 (one mismatch allowed), −1 = 15 (15 bp seed), −y = try hard, and −m = 1.

We detected 66,893 unique TA insertion sites in our input library. We observed a significant bottleneck in all plant and clay treatments corresponding to an average loss of 38%, 35%, and 33% of the insertions in the Col-0, *deps*, and clay samples respectively. The MaGenTA fitness calculations pool all insertions per gene before calculating fitness. Using this approach, we found that all but 192 (3%) of the genes with insertions in the in the input retained insertions in the Col-0, *deps* and clay samples.

Genes were considered to significantly affect fitness if insertions in them resulted in an average log_2_(5)-fold increase or decrease in fitness and they had a p-value < 0.05 (Supplementary Data). To further study genes with large differential fitness between Col-0 and the *deps* rhizospheres, we looked at just genes that had a greater than −log_2_(10)-fold difference between the Col-0 and *deps* fitness scores once each was normalized to the clay-only control. We only considered genes with a normalized (rhizosphere / clay) fitness score <−log_2_(3) for Col-0 and >log_2_(3) for the *deps* mutant.

### Strain construction

Primers used for site-directed mutagenesis are listed in Supplementary Data.

Deletion strains *Pseudomonas* sp. WCS365 Δ*morA*, Δ*katB*, Δ*colR*, Δ*wapA*, Δ*gtsB*, and Δ*uvrA* were constructed by amplifying 500-700 bp of the upstream and downstream regions flanking the open reading frame and using overlap extension PCR (50) to join the two pieces prior to ligation into the pEXG2 vector (51). The pEXG2 vector confers gentamicin resistance and contains the *sacB* gene for counter-selection on sucrose. After confirming the correct insertion by sequencing, the plasmid was transformed into SM10λpir. Conjugations were performed with *Pseudomonas* sp. WCS365 by mixing a 2:1 ratio of washed overnight cultures of WCS365: SM10λpir with the desired plasmid, spotting onto King’s B plates, allowing the mating spots to dry, and incubating for 4 hours at 28°C. Mating mixes were then scraped off the plates and plated on selective media with 10 μg/mL gentamicin and 15 μg/mL. Successful integration of the plasmid into the genome confirmed by patching candidate colonies on sucrose or gentamicin. A second crossover event was selected by growing colonies overnight in media without selection and then plating sucrose without antibiotics. Candidate colonies were screened using primers outside of the initial construct and the final construct was confirmed by sequencing.

Deletion strains *Pseudomonas* sp. WCS365 Δ*spuC* and Δ*cioA* were constructed using a three-way cloning strategy. First, flanking regions of each gene were amplified using primers with added terminal restriction sites. The exterior ends of the regions were each modified with a unique restriction site, whereas a third restriction site was used for the interior ends of both regions. The suicide vector (pNPTS138) was then digested using the enzymes for the exterior ends, and each region was digested using the two enzymes appropriate for its own modified ends. The digested vector and the two flanking regions were then ligated and transformed into *E. coli*, followed by plasmid isolation and sequencing to ensure the integrity of the inserted deletion allele. pNPTS138 is a suicide vector developed for use in the Alphaproteobacteria *Caulobacter crescentus* (M.R.K. Alley, unpublished). Because it has a ColE1 origin which is specific to the Enterobacteriaceae, it should function as a suicide vector in *Pseudomonas*, which we confirmed by performing conjugations with an empty vector. Conjugations were carried out by mixing 1 mL of wild-type WCS365 with 1 mL of the WM3064 *E. coli* DAP (diaminopimelic acid) auxotroph strain carrying a suicide vector and plating 10 μL of the washed and concentrated cell mixture on LB supplemented with 0.3 mM DAP. After 4-6 hours, the “mating spot” was resuspended in 1 mL of supplement-free LB and dilutions were plated on LB with 50 mg/L kanamycin (LB-kan). Kanamycin-resistant WCS365 clones were restreaked on LB-Kan to purify, then patched densely onto no-salt LB with 10% sucrose to grow overnight as a lawn, which we then restreaked for single colonies. This was necessary because the *sacB* locus present on pNPTS138 did not confer strong sucrose sensitivity. This may be due to low expression of *sacB* in *Pseudomonas*, as Rietsch *et al*., developed the suicide vector pEXG2 for *P. aeruginosa* by adding a strong promoter to drive *sacB* expression (51). Nevertheless, 5%-10% of the single colonies grown on sucrose media were kanamycin-sensitive, indicating that there may have been weak sucrose counterselection. These Kan^S^ colonies were screened using PCR with the exterior primers for the flanking regions to distinguish strains with the deleted allele from wild-type revertants.

Site-directed mutagenesis of *Pseudomonas* sp. WCS365 *morA* GGDEF domain (Fig. 5A) was performed by amplifying *Pseudomonas* sp. WCS365 *morA* with FL05 & FL06; FL07 & FL08 and joining the product by overlap extension (50). Similarly, the EAL domain was mutagenized by joining the product amplified by FL01 & FL02 and FL03 & FL04. The joined PCR product was digested and ligated to pEXG2 vector for integration the WCS365 (51). Genomic mutations were confirmed by Sanger sequencing. D928AE929A mutations were introduced to the GGDEF domain (*morA^GGAAF^*); E1059A mutation was introduced to the EAL domain (*morA^AAL^*). These mutations were designed to abolish the catalytic activities of the diguanylate cyclase and phosphodiesterase, respectively (52–55). Screening of colonies to identify those with the correct mutations was performed using SNAP primers (56) designed to amplify the wild-type or mutant alleles (Supplementary Data).

pSMC21 (*Ptac-GFP*) and pCH216 (*Ptac-mCherry*) were transformed into wild-type or mutant *Pseudomonas* sp. WCS365 strains by pelleting an overnight culture, washing with 300 mM sucrose, and electroporating at 2.5 kV, 200 Ohm, 25 μF. Transformants were selected on LB with 50 μg/mL kanamycin. pCH216 was generated from pSMC21 (57) by excising GFP via a partial digest with XbaI and PstI and replacing it with PCR-amplified mCherry ligated into the XbaI and PstI sites.

### Annotation of candidate genes

WCS365_04639 was annotated as a catalase gene. PaperBLAST (http://papers.genomics.lbl.gov/cgi-bin/litSearch.cgi) result suggested that its product was highly similar to the *Pseudomonas* sp. SWB25 protein KatB (90% identity, 100% coverage) and to the *P. aeruginosa* PAO1 protein KatB (81% identity, 95% coverage).

WCS365_05664 gene product is similar to the previously characterized *P. aeruginosa* PAO1 protein MorA, a known diguanylate cyclase/phosphodiesterase (68% identity, 99% coverage) (23, 24).

WCS365_00305 was originally annotated as a putative aminotransferase gene. BLAST results suggested that WCS365_00305 encoded an aspartate aminotransferase; however, the most similar gene product based on PaperBLAST was SpuC (encoded by PA0299), a putrescine aminotransferase in *P. aeruginosa* PAO1.

Based on annotation, WCS365_05132 encodes a UvrABC system protein A, consistent with protein BLAST and PaperBLAST results. Its homolog *uvrA* in *Escherichia coli* encodes a subunit of the ABC excinuclease responsible for the repair of UV-induced thymine dimer and other DNA damage (37, 38).

WCS365_04646 was annotated as a cytochrome bd-I ubiquinol oxidase subunit 1 gene, consistent with PaperBLAST results. This gene is highly similar to the previously characterized *P. aeruginosa* PAO1 *cioA* (79% identity, 98% coverage), which, together with the gene product of *cioB*, forms CIO, a cytochrome with low affinity to oxygen (39).

The WCS365_04136 gene product is highly similar to the predicted amino acid sequence of *Pseudomonas* sp. SBW25 *gtsB* (PFLU4845), with 97% identity and 100% coverage, which has been shown to function as a glucose permease subunit of a ATP-binding cassette transporter (40).

Nucleotide BLAST result suggested that WCS365_05599 encodes a Type 6 secretion system (T6SS)-dependent secreted Rhs protein. Rhs was known to be a contact-dependent toxin delivered to neighboring bacterial cells, causing growth inhibition (41). However, no T6SS-related genes, such as *vgrG* or *hcp* genes, were found in the same contig as WCS365_05599 (58). Hence, we surmise that WCS365_05599 encodes a distantly related, T6SS-independent, contact-dependent toxin WapA (41).

### Rhizosphere and *in vitro* bacterial growth and fitness assays

Bacterial growth in the rhizosphere was quantified by growing *Arabidopsis* in 48-well clear-bottom plates with the roots submerged in hydroponic media and the leaves separated by a Teflon mesh disk (9). Plants were inoculated with wild-type and/or mutant *Pseudomonas* sp. WCS365 strains containing plasmids pSMC21 (*pTac-GFP*) or pCH216 (*pTac-mCherry*) and reading bacterial fluorescence with a SpectraMax i3x fluorescence plate reader (Molecular Devices; 481/515 GFP; 577/608 mCherry) (9). Briefly, 9 mm sterile Teflon mesh disks (Macmaster Carr) were placed individually in 48-well tissue-culture treated plates (Falcon). Each well was filled with 300 μl ½× MS media + 2% sucrose, and a single sterilized *Arabidopsis* seed was placed at the center of each disk. The media was replaced with 270 μL ½× MS Media with no sucrose on day 10, and plants were inoculated with 30 μL bacteria at an OD_600_ = 0.0002 (OD_600_ final = 0.00002; ~1000 cells per well) on day 12. For fitness assays, 15 μL each of the wild-type (mCherry) and mutant (GFP) strain were added to each well. To estimate the final relative proportion of each a bacterial strain, standard curves relating fluorescence intensity to bacterial OD_600_ of each fluorophore in each mutant background were generated. The fluorescence signal for each plant was measured pre-inoculation and this background was subtracted from the final well readings. The standard curve was used to estimate CFUs of each bacterial strain per well. The fraction that each strain contributed to the total bacterial population was determined. Data are an average of at least 3 experiments per bacterial genotype with a minimum of 6 wells per bacterial strain per experiment.

Bacterial growth and fitness *in vitro* were measured with a SpectraMax i3x plate reader (Molecular Devices). Overnight cultures were diluted to an OD_600_ = 1 in 10 mM MgSO_4_. 3 μL of diluted culture was added to 97 μL LB (rich media), M9 + 30 mM succinate (minimal media), or root exudate (described below).

### *In vitro* bacterial growth and fitness

Bacterial growth curves were performed by using bacterial cultures grown overnight in LB and then pelleted, washed in 10 mM MgSO_4_ and diluted to an OD_600_ =1. 3 μL of the culture was mixed with 97 μL of growth media for a starting OD_600_ = 0.03. Bacteria were grown in rich media (LB), minimal media (M9 salts + 30 mM succinate) or root exudate (M9 salts + 0.7× root exudate as the sole carbon source). Bacteria growth was quantified by measuring OD_600_ on a Versamax (Molecular Devices) plate reader. Doubling times were calculated using the exponential growth stage for each experiment and data reported are the average of 3 biological replicates.

For bacterial growth in competition with wild-type, mutant strains expressing GFP were mixed in a 1:1 ratio with wild-type strains expressing mCherry. Red and green fluorescence as well as OD_600_ were measured for each well. Using a standard curve generated for each fluorophore for each mutant or wild-type strain, the approximate bacterial OD was calculated and plotted as mutant growth in competition with wild-type (9). For fitness measurements, the fraction of the well represented by each mutant relative to the total bacterial growth in the well was calculated. For each experiment 3-4 technical replicates were performed and each growth curve was repeated at least 3 times. Doubling times were calculated using the exponential growth stage for each experiment and data reported are the average of 3 biological replicates.

Root exudate was collected by growing plants in 48-well plates for 12 days in ½× MS media with 2% sucrose (9). The media was replaced with ½× MS media with no sucrose and collected 1 week later. Exudate was pooled from multiple wells from 4 plates (~200 plants). Final root exudate contains spent MS media as well as potentially trace amounts of sucrose left from the initial plant media.

### Plant growth promotion (PGP) assays

The OD_600_ of *Pseudomonas* sp. WCS365 overnight cultures was measured before the cells were spun down at 10,000 × g for 3 min and washed with 10 mM MgSO_4_. After washing, cells were resuspended and diluted to OD_600_ of 0.01 in 10 mM MgSO_4_. Five-day old *A. thaliana* seedlings on ½× MS plates were inoculated at the root tips with 1 μL of diluted cell suspension. Images of growth promotion plates were taken with an Epson V850 scanner. Root length was quantified using the “Measure” function in Image J and lateral roots were counted manually using the scanned images.

### Histochemical GUS staining

Seedlings were grown in 96 wells plates in 100 μL 1× MS Media with 2% sucrose as described (2). The media was changed after 7 days, and bacteria were added to a final OD_600_ = 0.002 (20 μL OD_600_ = 0.01 per 80 μL media). GUS staining solution was added 16 hours later and incubated at 2 hours at 37°C. The GUS solution was removed and seedlings were cleared with 95% ethanol overnight for imaging.

### Crystal violet biofilm assays

Biofilm assays were performed as previously described (59, 60). Briefly, overnight culture of *Pseudomonas* sp. WCS365 cells were spun down at 10,000 × g for 3 min and washed twice, resuspended, and diluted to OD_600_ of 0.1 in with M63 medium (1× M63 salt, 0.2% glucose, 0.5% casamino acids, and 1 mM MgSO_4_), M63-putrescine medium (1× M63 salt, 0.2% glucose, 0.5% casamino acids, 1 mM putrescine, and 1 mM MgSO_4_), or M63R (1× M63 salt, 0.4% L-arginine, and 1 mM MgSO_4_) when appropriate. One hundred microlitres of diluted cultures were incubated at 27°C for 17 h in non-tissue culture-treated 96-well plates (Falcon; Product No. 351177). After incubation, the plate was rinsed in distilled water twice before staining the biofilm with 125 μL of 0.1% crystal violet for 10 min. Excess crystal violet was washed three times in distilled water, and the plate was dried overnight before solubilizing the crystal violet in 125 μL of 30% acetic acid for 10 min before transferring to a new 96-well, flat bottom plate (VWR; Catalogue No. 10062-900) for spectrophotometric reading. Absorbance was measured at 550 nm. Background signals were measured from wells containing 30% acetic acid and were subtracted from the absorbance reading. All average absorbance signals were normalized against the wild-type values.

### Motility assays

Motility assays were performed as previously described (25, 61). Overnight cultures were spun down at 10,000 × g for 3 min, washed, resuspended, and diluted to OD_600_ of 1.0 in with M9S (1× M9 supplemented with 10 mM sodium succinate). For swimming motility, M9S 0.3% agar plates were inoculated by stabbing the plates with an inoculation needle dipped in the diluted culture without completely piercing the agar. Plates were incubated at 27°C for 65 h before imaging. For surfing motility, 0.3% agar M9S medium supplemented with 0.4% of citrus pectin (Alfa Aesar; Catalogue No. J61021-22) was used. Plates were inoculated with 1 μL of diluted culture and incubated at 27°C for 24 h before imaging.

## Data Availability

The WCS365 Whole Genome Shotgun project has been deposited at DDBJ/ENA/GenBank under the accession PHHS00000000. The version described in this paper is version PHHS01000000.

## Acknowledgements

This work was supported by an NSERC Discovery Grant (NSERC-RGPIN-2016-04121), Canada Foundation for Innovation and Canada Research Chair grants awarded to C.H.H. and previous awards from MGH Toteston & Fund for Medical Discovery Fellowship grant 2014A051303 and the Gordon and Betty Moore Foundation through Grant GBMF 2550.01 to the Life Sciences Research Foundation (LSRF). R.A.M. is supported by a Simons Foundation Grant to the LSRF. Z.X.L. was supported by an NSERC URSA. This work was initiated in the Laboratory of Frederick Ausubel in the Department of Molecular Biology at Massachusetts General Hospital with support from NIH R37 grant GM48707 and NSF grant MCB-0519898.

## Supplementary Figure and Table Legends

**Figure S1. Treatments and setup for the Tn-Seq experiment**. (A) Wildtype and mutant *Arabidopsis* were grown in a sterile clay mix and inoculated with 10^4^ CFU *Pseudomonas* sp. WCS365/plant. CFU/gram of root was measured one week later by homogenizing plant roots and plating to count CFUs. Levels designate significance by ANOVA and Tukey’s HSD. (B) Plants were grown in a sterile soil-like mix of 1:1:1 of calcine clay:sand:pearlite saturated with ½x MS media with no carbon source and inoculated with a transposon insertion library of *Pseudomonas* sp. WCS365. The no plant control was inoculated with 20 mM succinate to allow for bacterial growth. Soil cores or plants roots were harvested 1 week after inoculation. Cores or roots with attached soil were used for DNA isolation. (C) Final growth of bacteria in soil mix (no carbon) soil mix with succinate, or in the rhizosphere of Col-0 or the quadruple hormone mutant.

**Figure S2. Phylogenomic analysis places *Pseudomonas* sp. WCS365 within the *P. brassicacearum* subgroup of the *P. fluorescens* group**. The genome of *Pseudomonas* sp. WCS365 was sequenced and a phylogenomic tree containing other *Pseudomonas* spp. was generated using PhyloPhlAn (Segata *et al*., 2013).

**Figure S3. Library construction and frequency of insertions by gene in Tn-Seq input library**. (A) Primers are shown in Supplementary Data. Step 1: After DNA isolation, DNA was digested with MmeI to cleave 21 bp upstream and downstream of the transposon insertion. Step 2: Digested DNA was phosphotased. Step 3: Double stranded adapters were ligated onto phosphotased DNA. Step 4: The region flanking the transposon junctions was PCR amplified using a transposon-specific (PCR.X) and adapter specific (U.) primer. (B) The input library was sequenced and mapped to the WCS365 genome. We found a mean of 10.3 and median of 8 insertions per gene.

**Figure S4. Growth of *Pseudomonas* sp. WCS365 mutants in rich and minimal media and the Δ*gtsB* mutant in dextrose**. (A-C) Growth of *Pseudomonas* sp. WCS365 and mutants in (A) LB, (B) M9 + 30 mM succinate, and (C) M9 + root exudate. For all assays, *Pseudomonas* sp. WCS365 were grown overnight and then diluted to an estimated OD_600_ = 0.03. Bacteria were grown for 24 hours in a shaking plate reader with readings taken every 15 minutes. (D-F) The Δ*gtsB* mutant was growth in (A) LB, (B) M9+30 mM succinate, or (C) M9+20 mM dextrose.

**Figure S5. Images of *Pseudomonas* sp. WCS365 plant growth promotion (PGP) assays**. Plants were grown on plates and inoculated with wildtype or mutant *Pseudomonas* sp. WCS365 with 3μl bacteria at a final OD_600_ of 0.01. Plates were imaged 10 days later and lateral root density was calculated and shown in Fig. 3.

**Figure S6. Biofilm and motility phenotypes of *Pseudomonas* sp. WCS365 mutants**. (A) Swarming motility and (B) surfing motility assays with WCS365 mutants. All data were normalized for the wildtype control of a given experiment. (C-D) Crystal Violet Assays were performed in (C) minimal M63 media or (D) M63 supplemented with arginine. (A-B) Data are the average of 3 biological replicates with 3 technical replicates per experiment; \**p*<0.05, \*\**p*<0.01 by ANOVA and Tukey’s HSD. (C-D) Data shown are the average of 3 biological replicates with 8 technical replicates per experiment; *and ** *p*<0.05 by ANOVA and Tukey’s HSD.

**Figure S7. The *Pseudomonas* sp. WCS365 Δ*spuC* mutant cannot use putrescine as a sole carbon source**. (A) Growth of wild-type and the Δ*spuC* mutant in M9 + 25 mM succinate or (B) M9 + 25 mM putrescine.

**Table S1. Doubling times for *Pseudomonas* sp. WCS365 mutants *in planta* and *in vitro***. Mutants were grown in LB, M9 + 30 mM succinate, or M9 with root exudate as the sole carbon source. Bacterial OD_600_ was measured with a plate reader (Fig. S5) and doubling time was calculated based on growth during exponential stage. Relative fitness was calculated as the fraction of the final culture that was the designated mutant strain when grown in competition with wild-type. The rhizosphere data are shown in Fig. 2a, and the in LB, M9 succinate and M9 root exudate were calculated based on 20 hours of *in vitro* growth and shown in Fig. 2C-E. \**p*<0.05 and \*\**p*<0.01 by ANOVA and Tukey’s HSD relative to wild-type *Pseudomonas* sp. WCS365.

**Table S2. Fitness of *Pseudomonas* sp. WCS365 mutants *in planta* and *in vitro***. Relative fitness was calculated as the fraction of the final culture that was the designated mutant strain when grown in competition with wildtype. The rhizosphere data are shown in Fig. 2A, and the in LB, M9 succinate and M9 root exudate were calculated based on 20 hours of *in vitro* growth and shown in Fig. 2C-E. \**p*<0.05 and \*\**p*<0.01 by ANOVA and Tukey’s HSD relative to wildtype *Pseudomonas* sp. WCS365.

